# Long-term cold selection leads to increased thermal plasticity of male body size and thermal canalization of fecundity in *Drosophila melanogaster*

**DOI:** 10.1101/2023.11.13.566815

**Authors:** Rishav Roy, Aradhya Chattopadhyay, Sreebes Deb Sharma, Aharna Mondal, Payel Biswas, Shampa M. Ghosh

**Affiliations:** School of Biotechnology, Kalinga Institute of Industrial Technology (KIIT), Bhubaneswar, India; School of Biological Sciences, University of Edinburgh, UK

**Author notes:** Joint first authors (Rishav Roy, Aradhya Chattopadhyay).

**Keywords:** Thermal plasticity, Thermal adaptation, Experimental evolution, Fecundity, Body size, Canalization

## Abstract

From laboratory populations of *Drosophila melanogaster* maintained at 25°C for many decades, new populations were derived and maintained at 17°C for over 60 generations. Fitness traits such as body size and fecundity of the cold and warm selected populations were quantified at both 17°C and 25°C. Flies from both selection regimes had similar body weight when grown at 25°C, but 17°C selected males had greater dry weight and less relative water content than 25°C selected males, when grown at 17°C. Cold selected males thus evolved greater thermal plasticity of body weight and density compared to controls. Fecundity data showed, 25°C selected females laid less eggs at 17°C compared to 25°C treatment, but 17°C selected females did not show significant difference in fecundity between 25°C and 17°C. Cold evolved flies thus showed thermal canalization of fecundity, possibly buffering the repressing effect of cold. Whether or not (i) greater body mass and density of cold evolved males, and (b) unrepressed fecundity of cold evolved females, both observed when reared at cold, are causally related, remain to be investigated. The study thus shows thermal selection can lead to evolution of greater thermal plasticity in a trait, while result in thermal canalization of another.

## Introduction

Temperature impacts a wide array of biological traits, starting from molecules to cellular attributes, from physiology to development, and from behavioural to life history traits. Ectothermic organisms experience a greater impact of temperature because they do not have internal thermoregulation the way endotherms do [reviewed in [1]]. In ectotherms such as insects, temperature can induce both plastic and evolutionary changes. The short-term or instantaneous effect of temperature on a trait, depicted as thermal plasticity, is observed in a wide variety of biological traits. Such plastic changes are viewed as inevitable effect of temperature on biological parameters, like growth, metabolism, physiology etc.[2–5]. However, there are ample empirical evidences that suggest there could be genetic variation within and across species controlling how a trait is influenced by temperature, and hence thermal plasticity can be shaped by selection [6–10]. In addition, temperature can lead to evolution of traits in the long run, manifesting thermal evolution, or thermal adaptation [8,10,11]. Thus, temperature induces both (a) proximate changes or plasticity and (b) ultimate changes or evolution, in biological traits. However, how thermal plasticity and thermal evolution impact each other is not fully understood [8,10,12–14].

*Drosophila,* the ectotherm is particularly well-suited for studies focusing on both thermal plasticity and evolution, because of its (a) amenability to studies manipulating temperature in the laboratory, both within and across generations, and, (b) wide geographic range across latitudes and continents covering thermally diverse regions [15–17]. In holometabolous insects like *Drosophila*, cold developmental temperature leads to slow growth and metamorphosis, which translates into emergence of bigger flies, demonstrating the temperature size rule [5,18,19]. Cold temperature during the adult stage usually leads to prolonged lifespan but reduced reproductive output or fecundity [20,21]. Such kind of plastic effect of cold temperature on growth, body size and fecundity have been found to be consistent across studies, and is well documented in fly literature. However, long-term evolutionary effect of temperature on these traits may vary from population to population, as it seems to depend upon genetic composition and evolutionary history of the concerned population, and also upon specific thermal range considered for the study.

The objective of this study is to obtain a clear understanding of the relationships between temperature, fitness, plasticity, and adaptation in *Drosophila melanogaster*, using a well-defined experimental design. In view of these, we investigated how temperature shapes fitness traits like fecundity and body weight in *Drosophila*, at both plastic and evolutionary levels. Studies exploring the relationship between temperature and fitness have led to findings and given rise to hypotheses that are as diverse as the species and populations under investigation, their thermal histories, temperature ranges, and study designs. Some of these studies lend support to the optimal developmental temperature hypothesis, which suggests an organism achieves its highest fitness when it is raised at an optimal intermediate temperature [22,23]. Another hypothesis, namely the beneficial acclimation or the adaptive acclimation hypothesis states that individuals acclimated to a certain environment (*e.g.,* a temperature) perform better in that environment compared to individuals acclimated to a different environment [22,24,25]. Some studies also suggest adult fitness is greatest when the ambient temperature of the adult matches with the ambient temperature of the pre-adult stage [26]. There are two other hypotheses connecting temperature and fitness in various organisms. The colder-is-better hypothesis predicts individuals reared at low temperatures will have greater fitness than those reared at high temperatures; while the hotter-is-better hypothesis predicts the opposite [27–29]. Despite the studies and hypotheses, a tangible gap persists in the comprehensive understanding of the links connecting temperature, adaptation and plasticity and fitness.

Studies performed on fly populations collected from different latitudes and thermal ranges have added important information about thermal adaptation, but these studies have their own limitations. Evolution of latitudinal cline of flies may be influenced by factors other than temperature, like humidity, photoperiod, and altitude, among others. Therefore, some of the evolved changes may have been shaped by multidimensional selection pressures including factors additional to temperature. An alternative approach allows one to study adaptive evolution under a less complex scenario, and a more straight-forward manner. The *laboratory selection* or *experimental evolution* approach enables one to study evolution in carefully controlled set up of the laboratory, under the influence of clearly defined selection pressures. Studies employing experimental evolution have undergone much refinement in the last 50 years, and have enriched our understanding of adaptive evolution in an unforeseen manner [30–34]. While experimental evolution studies may not mimic multidimensional and complex selection pressures experienced by populations in the natural environment, these studies nonetheless hold considerable merit in delineating evolutionary responses of different selection pressures. Consequently, tracking evolution of populations subjected to specific thermal regimes under regulated laboratory conditions for many generations, can potentially shed light on the unique role temperature plays in shaping adaptive evolution and fitness.

Studies done in the past have employed laboratory selection or experimental evolution as an approach to study thermal selection [11,20,35–44]. Some of these studies were conducted for a very short duration [35,36], while some employed a small population size, making it difficult to make meaningful conclusions at the evolutionary level [37]. Some selection studies focused on comparing responses to constant *vs.* fluctuating thermal conditions [38,39], while others subjected large fly populations to evolve at different constant temperatures. One long-term thermal selection study that involved extensive characterization of life history, growth and size traits was conducted in Linda Partridge’s laboratory. Partridge group allowed populations of *D. melanogaster* to evolve at warm (25°C) and cold temperatures (first et 18°C, then gradually brought to 16.5°C) for many years [11,20,41].The populations adapted to warmer conditions exhibited higher fitness levels when raised in warm temperatures, while populations adapted to colder conditions performed better when exposed to colder temperatures [20], lending support to the adaptive acclimation hypothesis. Perhaps the study design of Partridge group matched closely with what fly populations might be facing while evolving in warm *vs.* cold environments in nature, however, similar to clinal studies, selection pressures experienced by these populations might have included factors other than that of temperature. To be more precise, pre-adult and adult population densities were not controlled in this study, and warm *vs.* cold rearing might result in very different population densities, which could potentially add some inadvertent selection pressures on body size and fitness through factors other than temperature, like crowding. Moreover, the populations were under overlapping generations, which can potentially result in age-structuring and reduction of effective population size, and these could be very different across warm and cold conditions. Given these limitations, evolution of these populations cannot be completely attributed to thermal selection, as confounding effects of crowding, age-structuring and dissimilar effective population sizes cannot be completely ruled out. Another study was conducted by research group of Mauro Santos, which involved laboratory selection of populations of *D. subobscura* at constant temperatures of 13°C, 18°C, and 22°C for over 4 years. They used discrete generation cycle and controlled larval and adult densities. These populations diverged for many traits including wing size and shape, development time, viability, gene expression, chromosomal traits [42,43]. Warm adapted populations (22°C) showed greater net fitness at all three temperatures (13°C, 18°C, and 22°C), while cold adapted populations had very low fitness in the warm environment (22°C) [44].

We performed a long-term thermal selection experiment using populations of *D. melanogaster*, with a rigorous experimental design that employed controlled rearing densities and discrete generations, among other factors. We used large outbred laboratory populations of *D. melanogaster*, which have been reared at the standard rearing temperature of 25°C for decades with rigorously controlled rearing densities and discrete generation cycles, and another set of populations, genetically related to the former, which have been selected with similar rigour at a colder temperature of 17°C, for close to 60 generations. Thermal range of *D. melanogaster* ranges from ∼11°C to ∼32°C [17,23]. While 25°C is optimal for these flies and can be considered warm, 17°C is sufficiently cold, which lengthens the development time two-fold compared to that at 25°C, produces much bigger flies [5,7], and suppresses fecundity [20,21]. We aimed to see when subjected to evolve at 17°C for many generations, how fitness traits evolve and diverge in these populations, compared to the ancestral controls maintained at 25°C. Reproductive output is a direct measure of Darwinian fitness of an organism, and in *Drosophila*, it can be measured as either the number of eggs laid, or the offsprings produced by the flies. We used fly fecundity, *i. e.* the number of eggs laid per female as a measure of reproductive output or fitness of flies. Another trait, which is an indicator of fitness is body size of flies. Body size is positively correlated with (a) female fecundity and (b) male mating success in *Drosophila* [45,46],reviewed in [31] Therefore, we investigated the evolved and plastic changes in the body size of flies caused by temperature.

To quantify reproductive output, or fecundity, we measured eggs laid by individual females every day, up to 22 days of adult life. *Drosophila melanogaster* lives for about 35-40 days on average, at standard rearing temperature of 25°C, and the female lays maximum number of eggs in early 3-4 days of life, after which the fecundity dwindles for the fly [reviewed in [31]. Therefore, 22 days fecundity is a fairly good representation of the overall egg output of the fly. We employed a 2 × 2 full factorial design in which flies from the two thermal selection regimes were grown and assayed at two treatment temperatures. This design allowed investigation of the joint effects of selection and treatment temperature on fecundity.

In majority of studies focusing on body size in *Drosophila*, wing size has been used as a proxy for body size. This is common in studies investigating body size clines, cellular bases of size, etc[17,24]. However, some studies have also used thorax size [9,47], while others have used body weight as a measure of body size in flies [reviewed in [31] While there are some suggestions that for a tiny insect like *Drosophila* a linear measurement like wing, thorax or leg might be a better indicator of body size than body mass [47], individual body parts may show different degree of sensitivity to environmental factors like temperature. For example, wing size is highly sensitive to rearing temperature while thorax and leg size show moderate and low sensitivity to temperature [48],Ghosh and Shingleton, *unpublished data*], and therefore, thorax and leg size may not be a reliable indicator of body size variation generated by temperature. On the other hand, tiny size and very low weight of flies notwithstanding, body weight, wing size and body size have been found to be positively correlated in flies [15] and body weight has been used frequently as a measure of body size in studies investigating life history evolution [reviewed in [37]]. Moreover, in a recent study, authors found a strong positive correlation between wing size and body weight, across 38 strains of the *Drosophila* Genetic Reference Panel or DGRP [49]. Therefore, body weight can be reliably used as a proxy for body size in flies, and we used the same, but with some caution. There are some concerns that body weight of a fly can vary considerably depending upon its age, activity, and in case of females its egg laying status (pre and post peak fecundity). Taking the above factors into consideration, we measured weight of only freshly eclosed unmated flies, within 4 hours of emergence from pupae, keeping age, activity, and reproductive status consistent.

The 2 × 2 full factorial design used in our study allowed us to investigate both (a) long-term or evolutionary effect of cold *vs.* warm selection, and (b) short-term or plastic effect of warm and cold temperature on fitness traits like fecundity and body size of flies. While we measured egg production of flies for fecundity, we assayed both wet and dry body mass of flies for measuring body size. This also allowed us to find out relative content of water to body mass of the flies. It is also important to note, we investigated the *selection response* in fecundity and body weight caused by temperature by comparing the main effect of selection temperature on the traits. Comparing the change in the trait values across treatment temperatures for a given selection regime allowed us to evaluate the extent of *plasticity*, or the lack thereof (referred to as *canalization*), for these traits. The study design also enabled us to find out whether selection in warm *vs.* cold has changed the extent of plasticity or canalization of the traits under study.

## Materials and Methods

### Study populations and selection regimes

We used six large outbred laboratory populations of *D. melanogaster*: three populations subjected to evolve at 17°C (KIIT Base populations: KB17 1-3), and three control (baseline) populations maintained at 25°C (New KIIT Base populations: NKB25 1-3). Both KB17 and NKB25 populations were derived from MB (melanogaster Base) populations, originally from the laboratory of Amitabh Joshi [50]. MB populations are descendants of flies derived from natural populations of *D. melanogaster*[50]. MBs and their ancestors have been selected at 25°C for many decades. Five replicates of MB populations (MB 1-5), selected for 202 generations on cornmeal medium at 25°C, ∼90% relative humidity and constant light on a 21-day discrete generation cycle [51] were brought to our laboratory in the summer of 2017. All 5 replicates were then mixed and this population was named KB-mix (KIIT Base mixed). KB-mix population was maintained on a 21-day discrete generation cycle on cornmeal medium at 25°C, ∼80% relative humidity and constant light for 2 years. Subsequently, KB-mix was split into three replicates, namely KB1, KB2, and KB3. After one generation of rearing, each replicate of KB was further split into two populations, one maintained at 17°C, another maintained at 25°C. They were christened KB17 1-3 and KB25 1-3 respectively. The KB17 and KB25 populations were maintained on similar conditions as KB-mix, but at their respective selection temperature.

Unfortunately, after 2.5 years of selection, all 3 replicates of KB25 populations died due to BOD incubator malfunction, and the populations could not be rescued. However, selection was continued for KB17 selection lines and they were used in our study. As we did not have matched controls for our 17°C selected lines, we used an alternative approach. We brought flies ancestrally related to KB17 that is maintained at 25°C. Laboratory of Imroze Khan had fly populations that were derived by mixing the 5 MB populations (of Amitabh Joshi laboratory) [52] and later split into 5 new replicate populations, known as MIB 1-5. MIB populations were maintained under same conditions as MBs except for the fact they had a 15 day generation cycle. We brought MIB 1-3, started maintaining them on a 14 day generation cycle, and rechristened them New KB25 1-3 (NKB 1-3). Thus, both our thermal selection lines, *i. e.* KB17s and NKB25s are maintained on a discrete generation cycle, ∼80% relative humidity, constant light, on cornmeal food.

KB17 1-3 and NKB25 1-3 have two notable contrasts. First, the former is maintained at 17°C whereas the latter is maintained at 25°C. The second difference lies in their generation time. *D. melanogaster* takes 17-20 days to develop from egg to adult at 17°C, whereas it takes 8-10 days to develop at 25°C. The generation time, therefore, is much longer at 17°C. Consequently, NKB25s are maintained on a 14 day generation cycle whereas KB17s are maintained on a 24 day generation cycle. Apart from this, similar maintenance protocols are followed for two selection regimes. For each replicate population, flies are grown in 25 vials containing food, and larval density is controlled at ∼70 per vial. Upon eclosion, flies from all 25 vials are transferred to plexiglass cages containing food smeared with supplementary live yeast-acetic acid paste. On 11^th^ day after previous generation egg collection in NKB25 populations, and 21^st^ day after previous egg collection in KB17 populations, eggs are collected for the next generation. Thus, egg collection is done on 4^th^ - 6^th^ day of adult age for NKB25, and 4^th^ - 7^th^ day of adult age for KB17 populations.

### Generation of flies for experiments

Body weight and fecundity of both 25°C and 17°C selection lines were assayed at two treatment temperatures, namely (a) 25°C, and (b) 17°C. For comparing the selection lines at (a) 25°C, flies from both 17°C and 25°C selection lines were reared at a common temperature of 25°C for one generation, and then their progeny were assayed at 25°C. This way both selection lines were standardized, and any non-genetic parental effect of divergent temperature on the progeny was eliminated, and only selection response or evolved changes between the two thermal selection lines could be identified. Similarly, to compare the selection response at 17°C, both 17°C and 25°C selected populations were reared at 17°C for one generation, and their progeny were subsequently assayed at 17°C. For any given treatment temperature, the flies were both grown and assayed at that temperature. The larval density was controlled at ∼70 per vial for generation of flies for all our experiments.

All assays reported here were conducted between 56-63 generations of selection for KB17 populations. The NKB25s were between 11 to 21 generations of selection during these assays; however, their ancestors, *i.e.* the MIB flies, MB flies, and their ancestors were maintained at 25°C-the standard laboratory temperature for *D. melanogaster* for decades. Hence, the number of generations is somewhat irrelevant for the NKB25s, and they should rather be considered as control or baseline populations that have been adapted to 25°C for very long.

### Fecundity assay

This assay was conducted after 56 generations of selection of KB17 populations. 20 vials of eggs were collected for all three replicates of KB17, over an oviposition window of 13-14 hours at 17°C, and the vials were transferred to 25°C. This step onwards the entire assay was conducted at 25°C. Upon eclosion, these flies were transferred to cages, and were referred to as standardized flies (for 25°C assay temperature). These flies were provided with food and excess yeast-acetic acid paste for 3 days, and their eggs were collected in vials over a small oviposition window of 3 hours. The flies growing from these eggs are referred to as assay flies. After 8 days of egg collection, flies started eclosing, and those eclosing during the first 4 hours were discarded, as they represent very early eclosing flies. From this point onwards virgin separation of freshly eclosed flies were done every 4 hours, till the next day, and adult males and females were collected in separate storage vials. Once flies were collected in sufficient numbers and covering complete middle part of the eclosion time distribution, virgin collection was stopped. Similar to very early flies, very late eclosed flies too were excluded from the assay. Given the NKB25 populations were already maintained at 25°C, no standardization was needed for them. Eggs were collected from them over an oviposition window of 3 hours and rest of the assay was conducted at 25°C. Once eclosed, these were our assay flies. Similar protocol was followed for virgin separation in these flies. Once virgin collection was completed for both selection lines, one male and one female from storage vials were introduced into one fecundity assay vial containing 2 mL of food. 15 such vials were set up for the fecundity assay for each selection regime and replicate population. The eggs laid in each vial by the female over a period of 24 h were counted under the microscope and the pair was transferred to a fresh vial containing food every day, and the process was repeated. If a male from a pair died, then it was replaced with an unmated male of the same age. If a female died, it was not replaced, and data from the same vial was included in the analysis up to the death of the female. Similarly, for treatment temperature of 17°C, eggs were collected directly from generation 56 flies of 17°C selection lines, and assay flies were generated at 17°C. Simultaneously, 25°C selected populations were standardized for a generation at 17°C, and fecundity assay was performed on their offsprings, also maintained at 17°C. Fecundity count was done for over 3 weeks of adult life, 22 days to be precise, for 25°C selected flies standardized at 17°C, and 17°C selected flies standardized at 25°C. 22 days total fecundity was analyzed using a 2 selection × 2 treatment temperature full factorial design. The total number of eggs laid by the female was averaged across 15 vials and used for the analysis. The fecundity data was also analyzed to investigate the age-specific pattern of egg production. For this, the data was split into 3 parts: first week’s fecundity, second week’s fecundity and third week’s fecundity. The data for day 22 was not included in the analysis.

### Weight assays

Dry and wet weights of flies from all replicates of NKB25 and KB17 populations were assayed after 63 generations of cold selection. The assays were conducted at both 25°C and 17°C, and flies generated through population standardization were assayed along with their control populations. Assay flies were collected within 4 hours of eclosion and stored immediately in microcentrifuge tubes (MCTs), and freeze-killed by keeping the flies at −20°C for 40-45 mins.

Wet weight was assayed for males and females separately. For each combination of selection regime, treatment temperature, replicate population, and sex, 8 MCTs were set up with 5 flies in each. After freeze-killing, these flies were taken out of the freezer and weighed immediately. 5 flies were weighed at a time. A Sartorius Quintix 35 (d = 0.01mg) balance was used for weighing the flies. Weight measured for 5 flies per MCT was divided by 5 to obtain the average weight per fly. After measuring the wet weight of flies, dry weight of the same flies were also assayed. For this, after collection of the wet weight data, flies were returned to their respective MCTs, and placed in a hot air oven set at 70°C. They were kept at 70°C for 36 hours, thus being dehydrated completely, and then taken out and weighed again.

### Relative water content of flies

Water content of flies was calculated by measuring the water lost during dehydration in the hot air oven. Total dry weight of 5 flies stored in each MCT was subtracted from the wet weight of the same flies to obtain the water content (in mg) of flies in each MCT. The relative water content was calculated by dividing the water content by wet weight of flies in the respective MCT (in percent).

### Statistical Analysis

Mixed model analyses of variance (ANOVA) were performed for all the traits, *i. e.* fecundity, wet weight, dry weight, and relative water content. Replicate population means were used for testing significance of fixed factors and their interactions in all analyses. Post hoc comparisons were done using Tukey’s honest significant difference (HSD) test. Data analyses were carried out on JMP Pro 17.

For analysis of fecundity data, 22 days total fecundity per female was averaged across vials to obtain replicate population means. ANOVA was performed on replicate population means. Selection and treatment temperature were treated as fixed factors, and replicate population was treated as random factor nested within selection and treatment temperature. We also analyzed week-wise fecundity data (1-7 days, 8-14 days, 15-21 days) to investigate the effects of selection and treatment temperature on early fecundity, mid-stage fecundity, and late fecundity till 21 days of adult life. A three-way ANOVA was conducted with selection, treatment, age (week post eclosion) as fixed factors, with the random factor population replicate nested within all three of these.

For both wet weight and dry weight measurements, mean weight per fly from each MCT was averaged across 8 MCTs to obtain the replicate population mean. ANOVA was performed on replicate population means of individual fly weight, with selection, treatment temperature, and sex being treated as fixed factors. Replicate population was treated as random factor nested within all 3 fixed factors. Similar analyses were performed separately for wet weight and dry weight data. The relative water content data were obtained in percentage, and therefore were subjected to arcsine-square root transformation to meet the normality assumption of ANOVA. Population means were used for testing significance of fixed factors such as selection, treatment temperature, sex, and their interactions.

## Results

### Total fecundity

After 56 generations of selection, KB17 selection lines did not show any significant difference in fecundity from NKB25 selection lines (table 1). Treatment had a significant main effect on fecundity (*p* = 0.0001) as it was significantly less at 17°C treatment compared to 25°C. However, the interaction of selection and treatment temperature was significant (*p* = 0.0356) (table 1). Post hoc comparisons showed NKB25 selection line had a significant drop in fecundity from 25°C to 17°C treatment temperature, but KB17 selection lines did not show any significant change between 25°C to 17°C treatment (figure 1 & 2).

**Figure 1:**
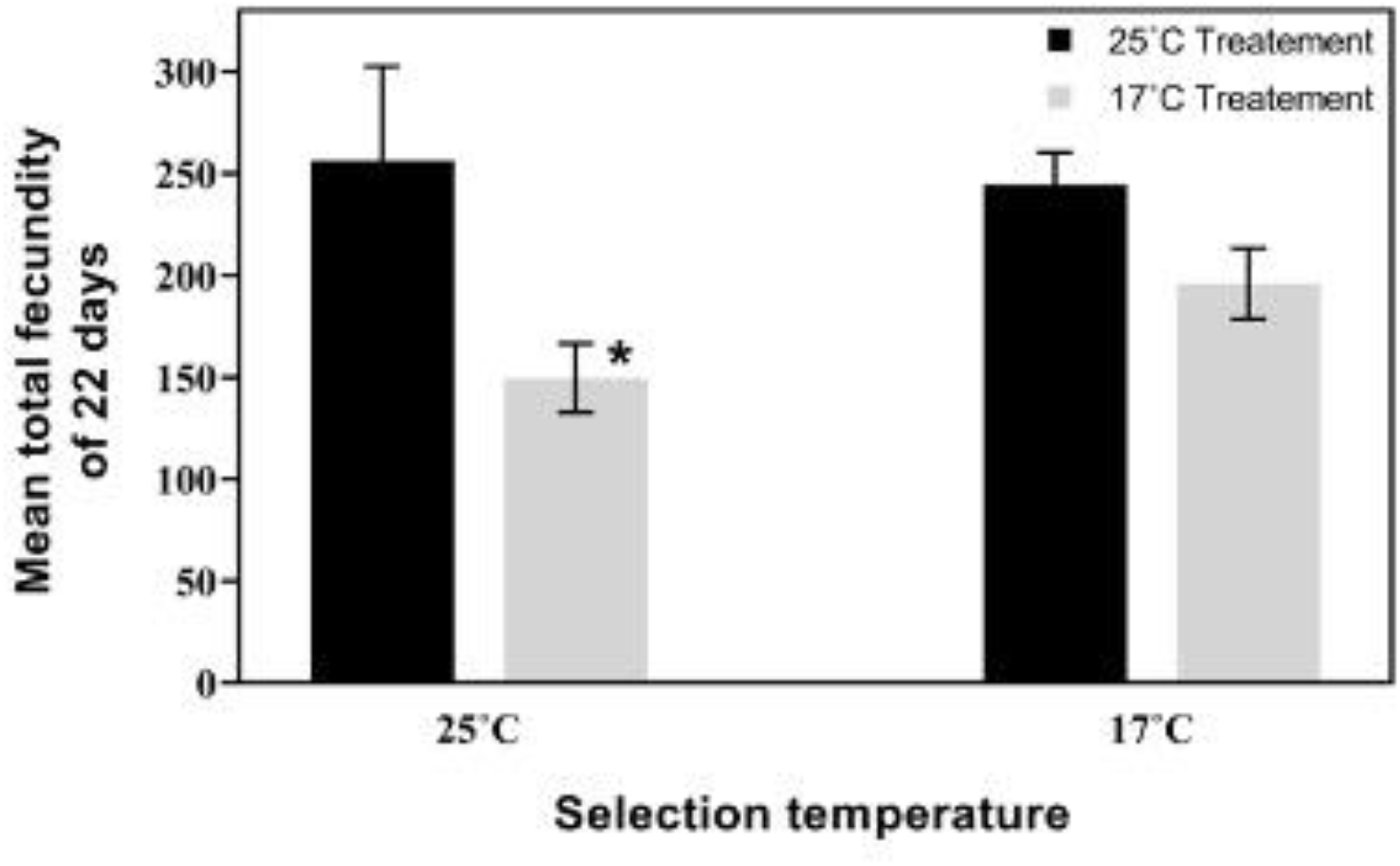
Mean total egg output per female over first 22 days of adult life in NKB25 and KB17 populations, assayed at 25°C and 17°C. The error bars represent standard error of the three replicate population means.

**Figure 2:**
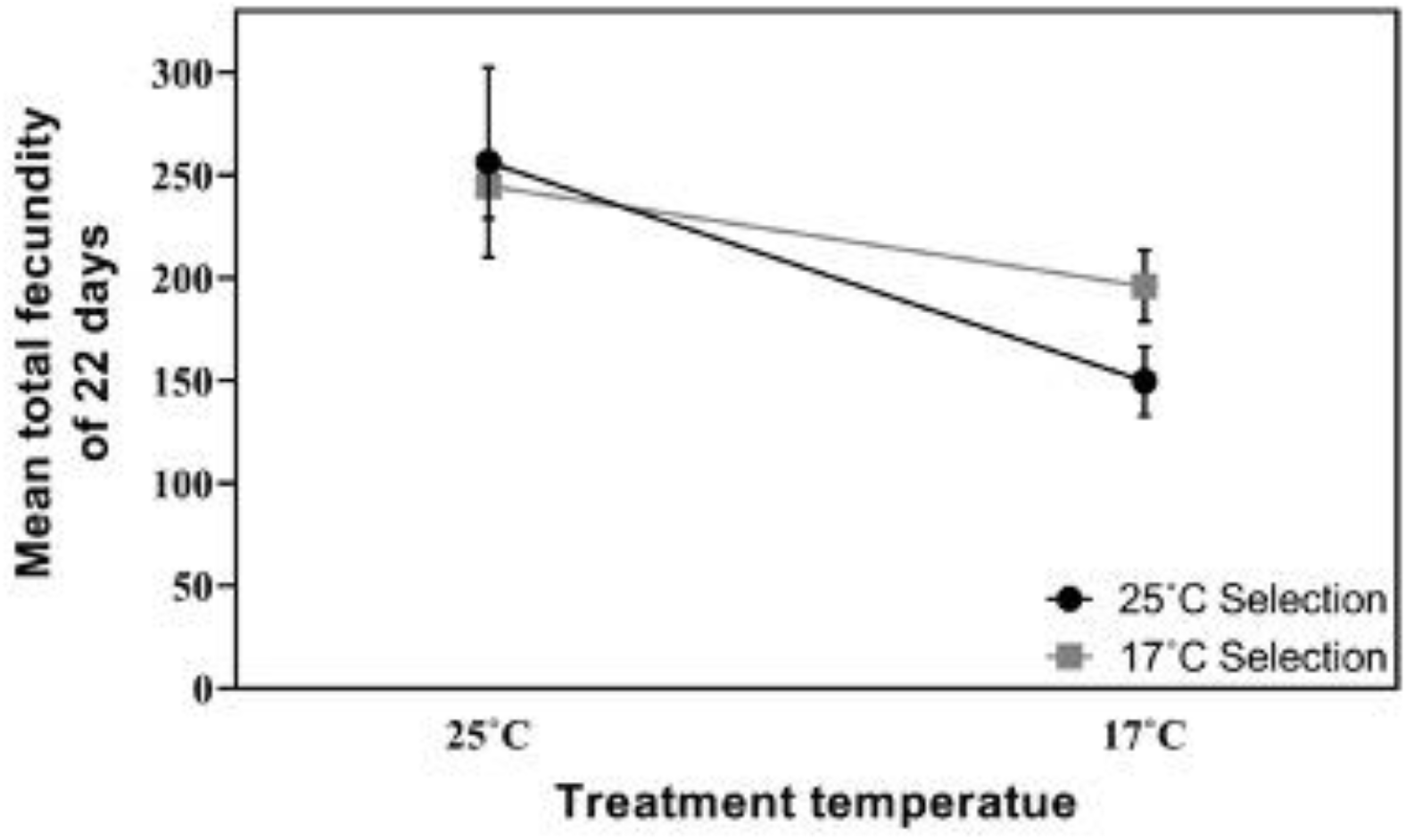
Reaction norms of mean total fecundity of 22 days of adult life in NKB25 and KB17 populations, reared at 25°C and 17°C. The error bars represent standard error of the three replicate population means.

**Table 1:** ANOVA results for mean total fecundity per pair for 1-22 days of adult life.

| Effect | *df* | MS | *F* | *P* |
| --- | --- | --- | --- | --- |
| Selection | 1 | 895.795 | 2.3083 | 0.1672 |
| Treatment | 1 | 17956.8 | 46.2714 | 0.0001 ** |
| Selection × Treatment | 1 | 2471.07 | 6.3675 | 0.0356 |

### Age-specific fecundity

Analysis of week-wise fecundity data showed significant effects of treatment, age, and interaction of treatment and age (table 2). 17°C treatment lowered overall fecundity, and fecundity was significantly higher during the first week compared to week 2 and 3 (figure 3). The treatment × age interaction showed fecundity was significantly higher during week 1 compared to 2 and 3 only at 25°C treatment, for both selection regimes. 17°C treatment did not show significant difference in fecundity across weeks. The main effect of selection and its interactions were not significant for the week-wise fecundity data. However, comparison of the means between section regimes showed an interesting trend. At 17°C treatment, fecundity was lower than 25°C overall, but KB17 had higher fecundity than NKB25 at 17°C, across all 3 weeks. First week’s fecundity at 17°C was 71% higher in KB17 compared to NKB25, while in the second week the same was 14% higher in KB17, and in the third week KB17 at 17°C laid 21% more eggs than NKB25s. 25°C treatment did not show difference in fecundity between NKB25 and KB17 during the first week, during second week, KB17 laid 9% less eggs than NK25, and in third week KB17 had 21% higher fecundity compared to NKB25.

**Figure 3:**
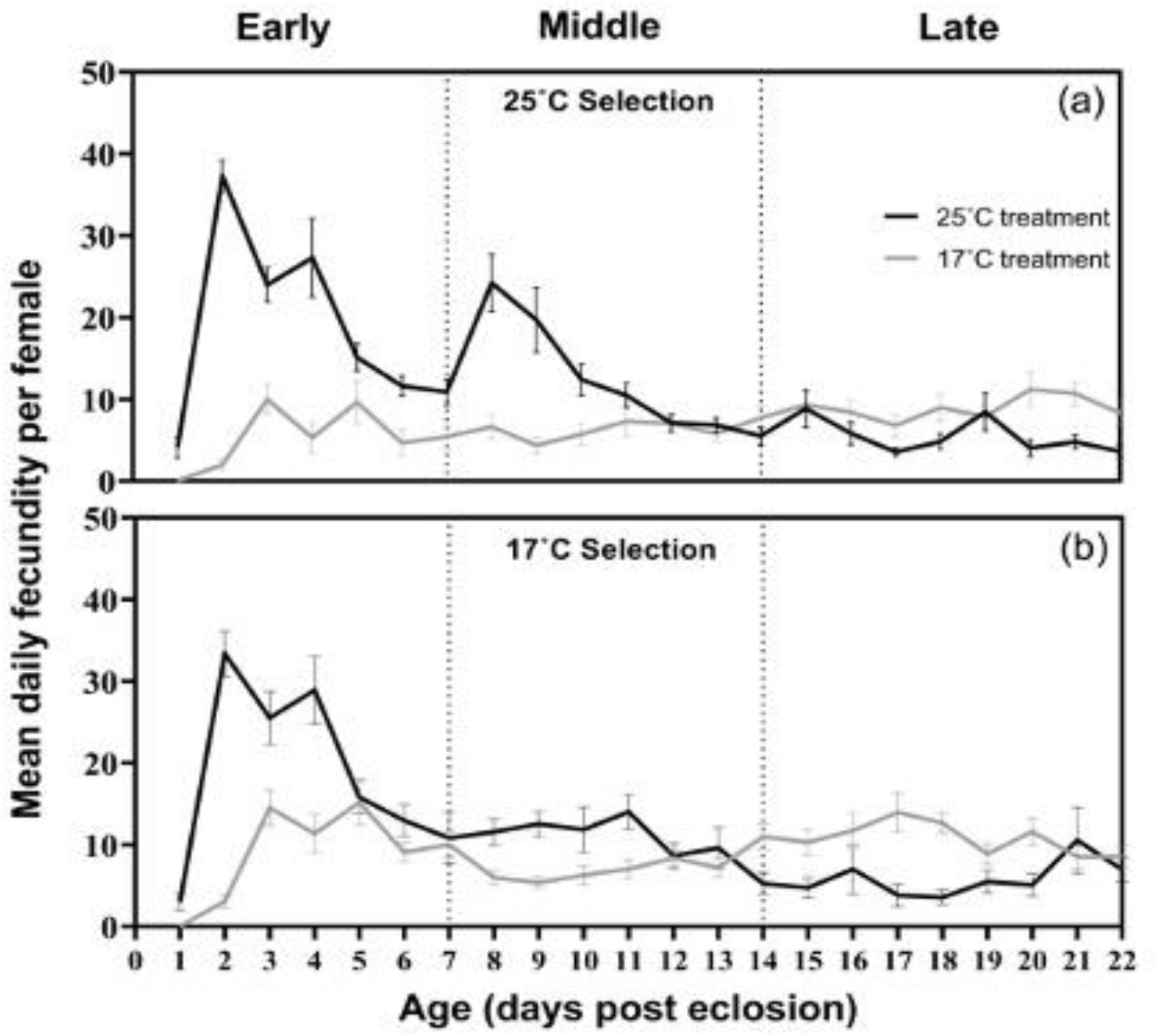
Mean daily egg output per female of (a) NKB25 and (b) KB17 populations over first 22 days of adult life, assayed at 17°C and 25°C. The error bars represent standard error of the three population replicate means. Early stands for day 1-7, mid for day 8-14, and late for day 15-22.

**Table 2:** ANOVA results for mean total fecundity per pair for early stage (1-7 days), middle stage (8-14 days), and late stage (15-22 days) of adult life.

| Effect | *df* | MS | *F* | *P* |
| --- | --- | --- | --- | --- |
| Selection | 1 | 543.849 | 2.5914 | 0.1205 |
| Treatment | 1 | 7844.29 | 37.377 | <.0001 |
| Week | 2 | 3525.06 | 16.7965 | <.0001 |
| Selection × Treatment | 1 | 527.684 | 2.5143 | 0.1259 |
| Selection × Age (week) | 2 | 166.809 | 0.7948 | 0.4632 |
| Treatment × Age (week) | 2 | 8445.4 | 40.2412 | <.0001 |
| Selection × Treatment × Age (week) | 2 | 82.1708 | 0.3915 | 0.6803 |

### Wet weight

After 63 generation of selection, wet weight of the flies was strongly affected by selection, treatment temperature and sex (figure 4, table 3). Overall, flies of KB17 selection lines were significantly heavier than their counterparts from NKB25 (*p* = 0.0084). Flies grown at 17°C had significantly greater wet weight than those grown at 25°C (*p* < 0.0001). Females had significantly greater wet weight than males (*p* < 0.0001). The two- and three-way interactions among selection, treatment temperature, and sex were not significant for wet weight. However, post hoc comparisons showed KB17 flies had significantly greater wet weight compared to NKB25 flies when grown at 17°C, but at 25°C both selection line flies had similar wet weights.

**Figure 4:**
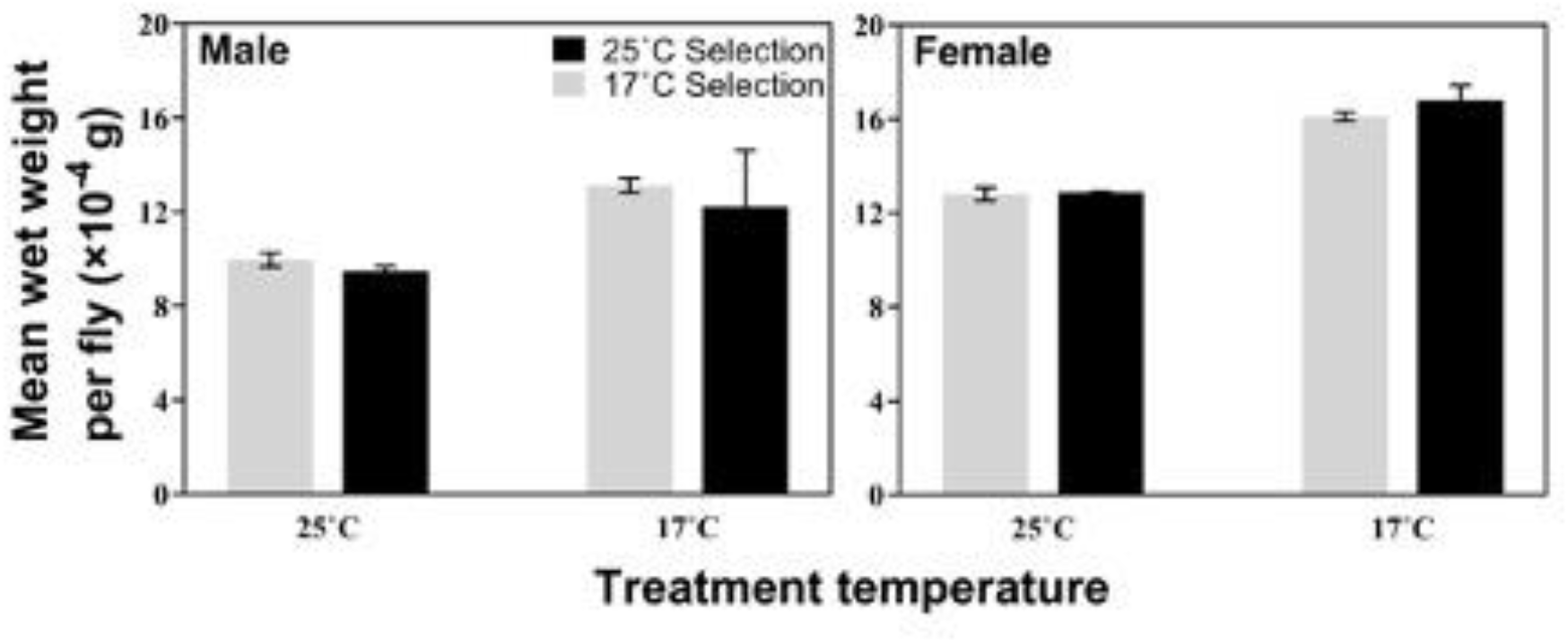
Mean wet weight per fly at eclosion in NKB25 and KB17 populations, reared at 25°C and 17°C. The error bars represent standard error of the three replicate population means.

**Table 3:** ANOVA results for mean wet weight per fly at eclosion Dry weight.

| Effect | *df* | *MS* | *F* | *P* |
| --- | --- | --- | --- | --- |
| Selection | 1 | 0.01735 | 9.0328 | 0.0084 |
| Treatment | 1 | 0.64329 | 334.9556 | <.0001 ** |
| Sex | 1 | 0.71544 | 372.5247 | <.0001 ** |
| Selection × Treatment | 1 | 0.00406 | 2.1153 | 0.1652 |
| Selection × Sex | 1 | 0.00132 | 0.6893 | 0.4186 |
| Treatment × Sex | 1 | 0.00686 | 3.5718 | 0.077 * |
| Selection × Treatment × Sex | 1 | 0.00013 | 0.0674 | 0.7984 |

Dry weight of the flies was strongly affected by selection, treatment temperature and sex (table 4). Overall, flies of KB17 selection lines were significantly heavier than NKB25 flies (*p* < 0.0001) (figure 5). Flies grown at 17°C had significantly greater dry weight than those grown at 25°C (*p* < 0.0001) (table 4). Females were significantly heavier than males across conditions (*p* <0.0001). Both selection × treatment, and selection × sex interactions were highly significant (*p* <0.0001). Dry weight of KB17 flies were significantly higher than NKB25 flies when grown at 17°C, but they were not significantly different from each other when grown at 25°C. Pooled over treatments, KB17 males had greater dry weight than NKB25 males, but the females from both regimes did not show significant difference in their dry weight overall. Treatment temperature × sex interaction was not significant. However, the three-way interaction among selection, treatment temperature and sex was highly significant (*p* = 0.0004). Post hoc comparisons showed KB17 males had significantly greater dry weight than NKB25 males when grown at 17°C, but their dry weight was not different when grown at 25°C (figure 5). Females from both selection regimes did not differ in dry weight at any of the two treatment temperatures. Interestingly, males at all combination of selection and treatment temperatures were significantly lighter than females, except for 17°C selection lines grown at 17°C, in which male and female dry weight had no difference.

**Figure 5:**
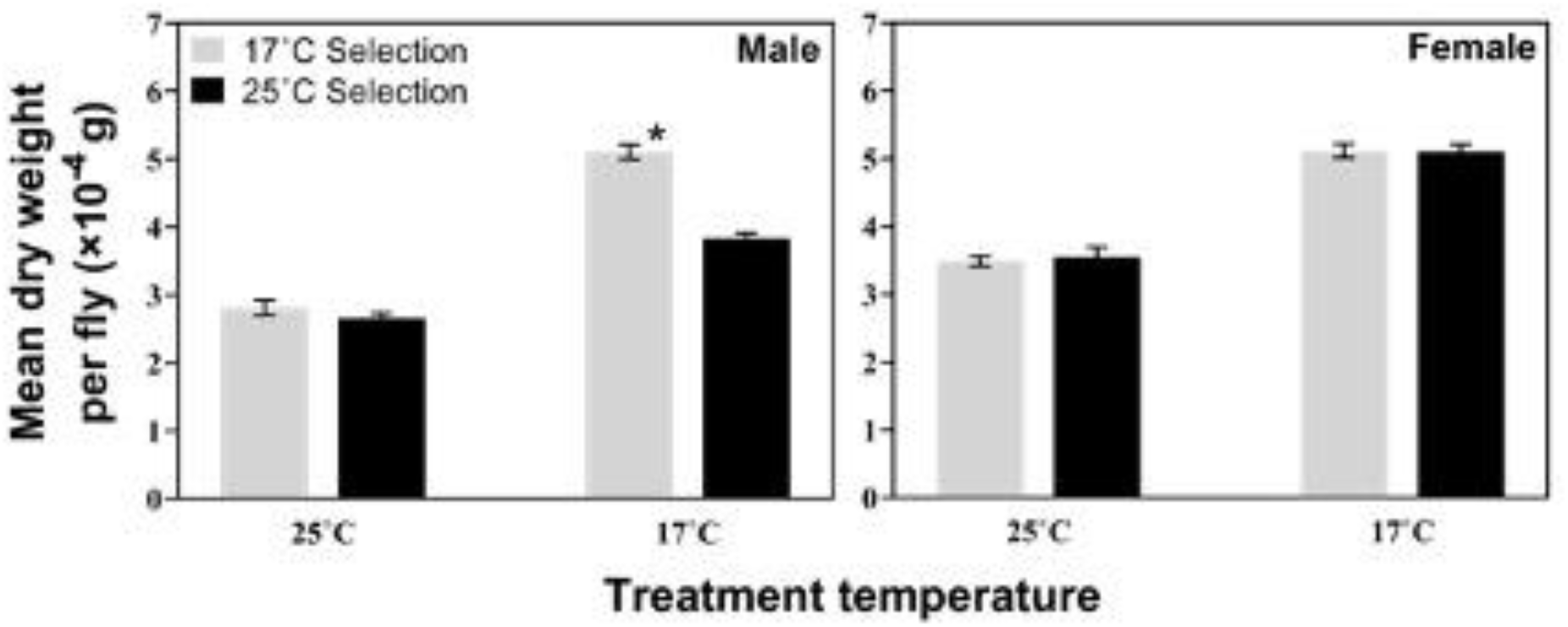
Mean dry weight per fly at eclosion in NKB25 and KB17 populations, reared at 25°C and 17°C. The error bars represent standard error of the three replicate population means.

**Table 4:** ANOVA results for mean dry weight per fly at eclosion.

| Effect | *df* | MS | *F* | *P* |
| --- | --- | --- | --- | --- |
| Selection | 1 | 0.00714 | 34.5274 | <.0001 ** |
| Treatment | 1 | 0.16581 | 802.208 | <.0001 ** |
| Sex | 1 | 0.03066 | 148.352 | <.0001 ** |
| Selection × Treatment | 1 | 0.0053 | 25.6407 | 0.0001 ** |
| Selection × Sex | 1 | 0.00812 | 39.2684 | <.0001 ** |
| Treatment × Sex | 1 | 0.00032 | 1.5384 | 0.2327 |
| Selection × Treatment × Sex | 1 | 0.00409 | 19.7673 | 0.0004 ** |

### Relative water content of flies

Relative water content of flies (water content scaled by wet weight) was significantly affected by selection (*p* = 0.0123), treatment temperature (*p* < 0.0001), and sex (*p* = 0.0004) (table 5). Relative water content per unit wet weight was significantly greater in NKB25 (70.3%) compared to KB17 flies (68.85%). Similarly, relative water content was significantly greater in flies grown at 25°C (72.3%) compared to those grown at 17°C (66.9%). Females had significantly greater relative water content (70.85%) compared to males (68.4%). All two- and three-way interactions among selection temperature treatment temperature and sex were significant (table 5). KB17 males had significantly less water content (61%) compared to NKB25 males (68.6%) when grown at 17°C, but the males from the two regimes had similar relative water content when grown at 25°C (figure 6, table 5). Females from the two selection regimes did not show difference in relative water content at any of the treatment temperature.

**Figure 6:**
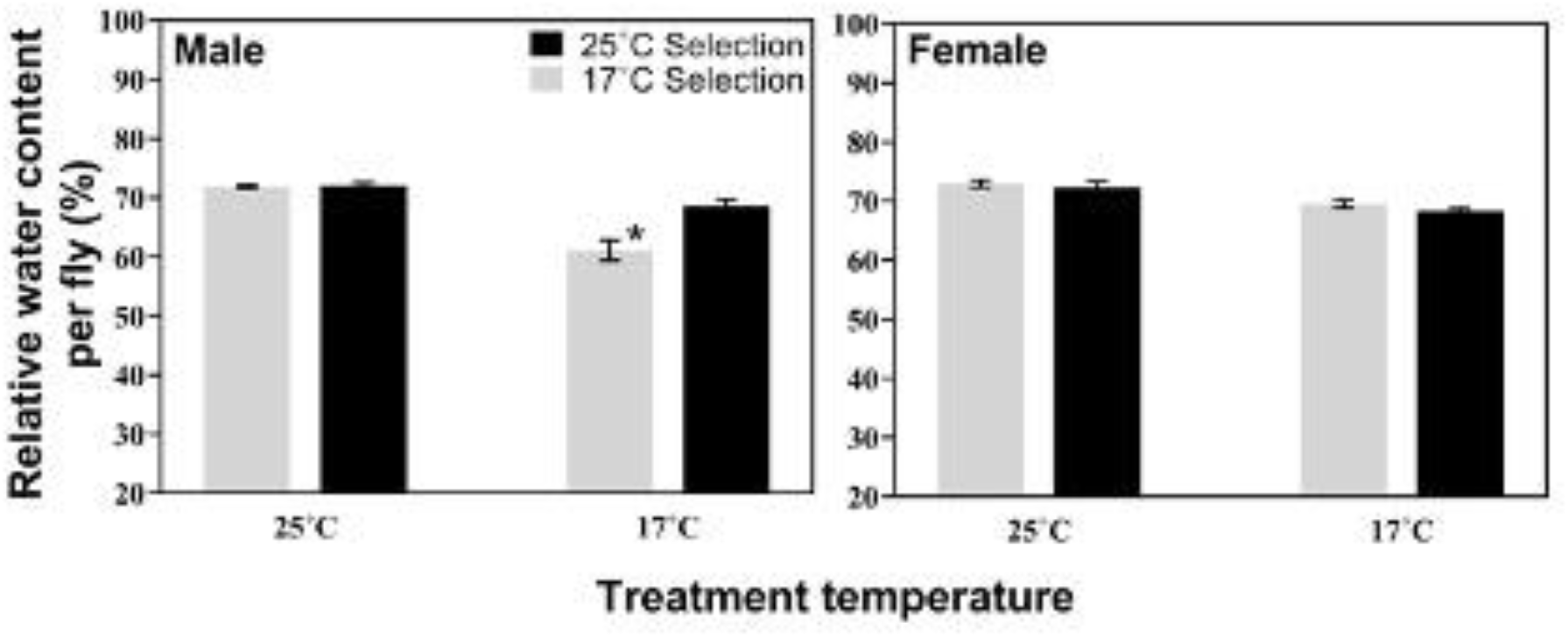
Mean relative water content per fly at eclosion in NKB25 and KB17 populations, reared at 25°C and 17°C. The error bars represent standard error of the three replicate population means.

**Table 5:** ANOVA results for mean water content scaled by wet weight per fly at eclosion.

| Effect | *df* | MS | *F* | *P* |
| --- | --- | --- | --- | --- |
| Selection | 1 | 13.2702 | 7.9574 | 0.0123 * |
| Treatment | 1 | 174.452 | 104.6097 | <.0001 ** |
| Sex | 1 | 33.9829 | 20.3777 | 0.0004 ** |
| Selection × Treatment | 1 | 17.2591 | 10.3494 | 0.0054 ** |
| Selection × Sex | 1 | 34.6921 | 20.803 | 0.0003 ** |
| Treatment × Sex | 1 | 17.748 | 10.6425 | 0.0049 ** |
| Selection × Treatment × Sex | 1 | 23.4481 | 14.0606 | 0.0017 ** |

## Discussion

### Evolution of the females: the fecundity

Cold temperature reduced pooled fecundity of the two selection regimes, but the interaction between selection and treatment revealed dissimilar effect of cold temperature on the warm and cold evolved populations. The control groups, *i. e.* the warm selection lines, suffered from a significant decline in fecundity when exposed to cold temperature, in comparison to their fecundity in warmer conditions. In contrast, the cold selection lines did not suffer such significant reduction in fecundity at cold treatment temperature compared to warm treatment. For *Drosophila melanogaster*, 25°C is known to be the optimal temperature, and findings from our control populations show cold temperature represses their fitness, as observed in numerous previous studies [21,23,26]. However, the cold selected populations seem to have evolved the ability to withstand cold, such that it does not affect their fecundity. This clearly indicates the cold evolved flies have adapted to cold as a result of selection. Surprisingly, cold adaptation did not lower their fitness in the warm environment as the fecundity of both selection lines were at par at the warm treatment. Our findings therefore did not reveal any trade-off between adaptation to 17°C *vs.* 25°C.

If we compare the reaction norm of fecundity levels across 17°C and 25°C treatments, 25°C adapted populations have a more plastic reaction norm while the one for 17°C selected populations is less plastic (figure 2). This means the thermal reaction norm of fecundity has evolved to be less plastic, or more canalized, in the cold selected populations. For a trait like fecundity, canalization, or robust reproductive output, across environments is likely to confer greater fitness and adaptability in diverse environments. The sustained fecundity of cold-adapted flies transitioning from warm to cold environments — an attribute not observed in warm-evolved controls, hints at an evolved mechanism neutralizing the cold’s suppressive effect. There could be different possible evolved mechanisms for the same. Some of the possibilities are as follows: (a) cold perception that potentially modulate egg production might have diverged in the two selection lines such that egg production of the cold evolved flies are not as affected by cold as the warm evolved flies; (b) cold selection might have led to bigger flies such that an increased size buffers against repressive effect of cold on fecundity, as body size and fecundity are positively correlated in flies. There could be possible alternative mechanisms accounting for the thermally canalized fecundity of cold evolved flies.

### Age-specific fecundity pattern

Apart from total fecundity, our experiment revealed notable impacts of selection and treatment temperature on the age-specific pattern of fecundity. The results showed several interesting trends for the pattern of fecundity. The distribution of fecundity along the age axis in wild-caught *Drosophila*, as well as those maintained in the laboratory tends to be positively skewed - a triangular shape of the lifetime fecundity distribution characterized by an early peak, is a typical feature of iteroparous insects [57]. This peak has been shown to be triggered by onset of mating in flies [58]. In our study, the typical early life spike in fecundity was observed at 25°C for both selection lines, which also coincides with the age of egg collection in our selection regimes. Surprisingly, we did not observe such a spike at 17°C treatment for either of the selection lines, which suggests the early life fecundity spike may not be canalized across conditions, and can be absent in colder environments.

The week-wise breakdown and analysis of fecundity data showed when flies were grown and assayed at 25°C, fecundity pooled across selection regimes was highest during first week, and then it showed a gradual decline during week 2 and 3. At 17°C, in contrast, the pooled fecundity did not differ across 3 weeks, rather the data show a somewhat moderate steady fecundity level at 17°C throughout the duration of the assay, which is different from the pattern observed at 25°C - the early spike followed by lower fecundity level afterwards. These contrasts between 25°C and 17°C treatments can be suitably explained by the lifespan of the flies at the said temperatures. Cold temperature increases lifespan in flies and a trade-off between lifespan and reproductive output is well-documented in evolutionary biology [reviewed in 31]. In conjunction with the existence of this trade-off, flies appear to maintain a relatively steady yet reduced egg production spanning their elongated lifespan at cold temperature. In contrast, at warm temperature flies lay maximum number of eggs early in life which subsequently dwindles to a lesser egg output for the remaining part of lifespan. The strategy at cold temperature seems to conserve resources, facilitating a prolonged lifespan and extended period of egg production. The energy needs to support a longer lifespan perhaps does not allow flies to blow up too much resources early in life. The age-specific fecundity pattern thus corresponds to energy need of the flies to live and reproduce for their respective lifespans at warm *vs.* cold conditions. It was observed that this pattern remained unaffected by selection.

Selection did not have a significant impact on week-wise fecundity data, but the data showed each week KB17 flies laid 14% to 71% more eggs on average compared to NKB25, at the cold treatment. This suggests, during each of the three weeks, low fecundity caused by cold treatment was much less prominent in KB17 populations compared to the controls, *i. e.* NKB25s. Perhaps variability of the fecundity data across individuals and replicate means did not result in a significant effect of selection on the week-wise egg output data, but like the total fecundity (of 22 days), these data showed the same trend, *i. e.* cold treatment temperature has less repressive effect on the cold selected populations. We speculate that in course of selection KB17 populations would show further improvement in their fecundity under cold condition. It is also important to note, the highest difference between KB17 and NKB25 fecundity at 17°C treatment was observed during the first week, when KB17 fecundity at 17°C was 71% higher on average than NKB25 fecundity. This is exactly what is expected, because the KB17 populations had been under selection for early life (day 3-4) fecundity at 17°C for 56 generations before this assay, and although early fecundity peak does not seem to be a feature of 17°C treatment, 17°C selection seemed to have been able to increase the early life fecundity at 17°C, showing signs of adaptation to the selection regime.

### Evolution of males: body size and relative water content at eclosion

Both dry and wet body weight of flies at eclosion were significantly influenced by treatment temperature, selection, and sex. Flies from the same selection regime were always heavier when grown at colder temperature. Selection temperature too had a significant main effect on body weight of flies, as cold evolved flies had both greater wet and dry weight than warm evolved flies. Females were heavier than males, under all conditions except one. The interactions among selection, treatment temperature, and sex, were more pronounced on dry weight compared to wet weight. The interactions showed that flies when grown at 17°C, had significant effect of selection, as the 17°C selected flies were heavier than 25°C selected flies. However, when grown at 25°C, the weights of flies from both thermal regimes were similar. Moreover, this effect was actually caused by the size divergence of males, as the 17°C selected males had 33% greater dry weight than 25°C selected males, when grown at 17°C. The weight of the females did not evolve in this study. Interestingly, as 17°C males evolved to have much greater dry weight when grown at 17°C while female size remain unchanged, the weight of the two sexes converged when grown at cold temperature. Consequently, sexual size dimorphism (SSD) that is ubiquitous in flies was absent in 17°C selected populations when grown at 17°C. In addition, analysis of relative water content of the flies showed 17°C selected males have also evolved significantly less relative water content than the 25°C selected males, when grown at 17°C. This indicates not only 17°C males evolved greater weight, but they were also more dense with relatively less water content, when grown at 17°C.

After 63 generations of evolving at cold, which took around 3.5 years, evolution of greater and more dense body mass only in male flies is really unique. Moreover, the existence of this difference only when the flies are grown at cold, but not at warm temperature indicates that the males of cold evolved populations undergo greater change in body mass and density between warm and cold rearing, compared to the males from warm evolved populations. This shows the cold evolved males have actually evolved *greater thermal plasticity of body mass and density*, a selection response not observed in the females. Given males and females share their genome, how the body mass or its plasticity have remained unchanged in cold evolved females is not clear yet. It is possible that the females may eventually evolve similar trend in course of selection, but, the body physiology and the selection pressure experienced by the two could be potentially different. Therefore, it would be erroneous to extrapolate findings from males to both sexes, and how the female body weight, physiology and plasticity may evolve as a result of cold adaptation may not be accurately predicted from evolution of the males. Nonetheless, the increased thermal plasticity of absolute and relative body mass of cold evolved males is an interesting evolutionary response of cold adaptation, and what is the underlying cause of this evolved change remains an interesting question to explore. Moreover, evolution of bigger male size that resulted in abolishing SSD in the cold evolved populations (when grown at cold) is a rare phenomenon, and how it affects the mating compatibility of the flies in these populations is a question relevant to sexual selection. Whether this is a transient phase of evolution of these populations, and if the females also eventually evolve bigger size or increased plasticity, remain to be observed in the coming years.

### Relationship between body size and fecundity

The coincidental increase in plasticity of male body weight and thermal canalization of female fecundity, caused by cold adaptation in our experimental evolution study are striking. Female body size is positively correlated with fecundity in a wide variety of organisms including *Drosophila* [reviewed in [31,46]. Had we observed an increase in female body size, or its plasticity in the cold evolved populations, better egg production of these flies in the cold, compared to that of the warm evolved flies could be attributed to the size advantage. However, the female size did not evolve in these lines yet, rather the males have evolved higher and more dense body mass. Male body size is positively correlated with lifetime mating success of *Drosophila* [45] as larger males are more successful at courtship and winning male-male competition for obtaining mates [53]. Larger males have also been shown to have higher postcopulatory success [54]. While a positive correlation between male body size and male reproductive success is clearly documented, it is less clear what effect male body size has on reproductive output of its female partners. It is generally observed larger males cause greater harm to the females by transferring more accessory gland proteins (Acp) during copulation, some of which are harmful for the females [55]. One study suggests smaller males might induce greater fecundity in their mates [45] while another study shows medium sized males lead to highest reproductive output in their mates [56]. However, the size variation of flies used in these studies were generated by manipulating larval densities, and hence reflect phenotypic plasticity of body size caused by the environment, *i. e.* pre-adult crowding. The size difference in male flies in our study is due to evolved or genetic difference in size caused by temperature. Moreover, in our study the females are not exposed to smaller *vs.* bigger males for the first time, rather they have co-evolved with the respective size of their mates. Therefore, how the size divergence of the males from the warm *vs.* cold selection lines may impact the fitness of their mates cannot be predicted from the documented effect of male size on female fitness where the size difference is neither genetic (*i. e.* evolved), nor caused by temperature. If two populations have evolved differences in male size, how it shapes the overall reproductive output of these two populations remains an open question. It is possible that in populations with larger average male body size, females overall end up laying more eggs on average due to possible greater reproductive potential of their mates. If this is true, then the cold-reared males of our cold evolved lines with their greater body mass, may induce greater fecundity in their mates. In our study, the cold and warm evolved female flies do not differ in size, yet the former are able to maintain their fecundity when grown at cold, unlike the warm evolved females who suffer reduced egg output at cold. This could be possible due to size advantage of the cold evolved males over warm evolved males when grown at cold, via their possible greater reproductive potential that might help the their mates (cold evolved females) to lay more eggs in cold, compared to those laid by warm evolved females. This would mean in cold evolved male size and populations, the higher thermal plasticity of body weight of males may help in greater canalization of female fecundity across temperatures. However, findings from this study are inadequate to prove this connection. Nonetheless, it does open up a pandora’s box linking male body size, female fecundity, thermal plasticity, and thermal evolution, and calls for a suite of rigorous experiments exploring these parameters.

### Findings and future direction

Our study shows cold adapted populations have greater fecundity than warm evolved populations when grown and assayed in cold environment, and in the warm environment, cold evolved flies are at par with the warm evolved flies. This shows although cold temperature like 17°C represses fecundity at the plastic level, long-term cold adaptation may to lead to evolution of greater fecundity, particularly in the cold environment. And this cold adaptation does not incur any cost on adaptation of cold evolved flies to the ancestral standard/warm environment. This study therefore, does not reveal any trade-off between adaptation to cold *vs.* warm environments as cold evolved flies in our study show greater fitness than warm evolved flies both in terms of male body weight and female fecundity, specifically at cold temperature, but they are at par with the warm adapted flies when grown at warm environment. However, it is important to note this study was conducted using a limited temperature range, confined to two set points (25°C as warm and 17°C as cold). Exploring extremes of the thermal spectrum, be it hotter or colder, might uncover additional aspects of adaptation to warm *vs.* cold, as suggested in previous research [17,22].

Our findings from the analyses of fecundity and weight data thus lead to several important findings about thermal evolution and thermal plasticity, as follows: (a) although cold temperature reduces reproductive output at the plastic level, cold adaptation can enable flies to overcome this suppression caused by cold. Despite adaptation to cold, the populations are not outperformed by the warm adapted populations when reared in warm environments. This suggests cold adaptation can thus lead to expansion of adaptive range of *D. melanogaster* populations rather than shifting their adaptive optima; (b) cold adaptation can increase the thermal plasticity of body weight and density, particularly in males, because at warm temperature warm and cold adapted males may not grow apart, but at cold temperature cold evolved males grow heavier and more dense than warm adapted males; (c) male size measured in terms of body weight at eclosion may evolve faster than that of females during cold adaptation. We suspect evolution of canalization of fecundity across warm and cold environments in cold evolved females can be possibly caused by at least in parts, greater thermal plasticity of body size of cold evolved males, which makes them bigger at cold, and may lead to greater fecundity of their mates at cold, such that the cold evolved females do not suffer a significant drop in fecundity; and (d) last, but not the least we show that the early life peak in fecundity oft-described in *Drosophila* may not be present at cold temperature, and flies would rather save on resources for living and reproducing for a longer time span at cold.

## Conclusion

Our findings show some rare empirical evidences of real-time evolution of a number of interesting phenomena due to cold adaptation, namely: (a) evolution of thermal canalization/ robustness of a fitness trait (fecundity), (b) evolution of greater plasticity of another fitness trait (body weight, and density), albeit observed only in males, (c) expansion of adaptive range of a population from warm to cold environment, and (d) evolution of bigger male size in absence of evolution of female size, leading to absence of SSD. The study also highlights and (d) absence of clear early life fecundity spike in *D. melanogaster* at cold temperature, typically observed at optimal temperatures like 25°C.

While our findings suggest cold adaptation in *D. melanogaster* may lead to expansion of thermal range of flies, and dismiss the presence of a trade-off in adaptation to warm *vs.* cold temperatures, it nonetheless signals the necessity of broadening the scope of the study. An ideal study should encompass a suite of other life-history traits and potentially extreme temperature conditions, to provide a holistic understanding of the adaptive capacities and potential trade-offs involved in thermal adaptation. Furthermore, it would be pertinent to delve deeper into the underlying mechanisms which account for the observed thermal canalization of fecundity in cold-adapted females, including the possible role played by the greater plasticity of the male body weight found in these populations. The conclusions drawn from this study, rooted in a rigorous 3.5 years of cold environment selection, certainly pave the way for intriguing questions about the future. There lies a captivating potential to observe the evolutionary ramifications unravelling in successive periods, looking 50 to 60 generations ahead, possibly unfolding a deeper layer of understanding of the evolutionary dynamics of thermal adaptation. The study lays a promising groundwork for further explorations into the rich tapestry of evolutionary biology, opening avenues for deeper inquiries into the nuanced relationships linking adaptation, canalization, plasticity, fitness, and selection.

## Acknwoledgements

We thank Amitabh Joshi, Imroze Khan and Abhiit Shit for providing flies from which our selection lines were initiated, Pragya Mishra, Proteek Sen and Subrath Ranjan for standardizing fly population maintenance and media preparation, Ranjit Pradhan and Manas Ranjan Mallik for general help in the laboratory.

## Author contributions

RR – Data curation, Formal analysis, Investigation, Methodology, Project administration, Resources, Supervision, Validation, Visualization, Writing-original draft, Writing-review and editing; AC – Data curation, Formal analysis, Investigation, Methodology, Project administration, Resources, Supervision, Validation, Visualization, Writing-original draft, Writing-review and editing; SDS – Data curation, Investigation, Methodology, Project administration; AM – Data curation, Investigation, Methodology, Project administration; PB – Data curation, Investigation, Methodology, Project administration; SG – Conceptualization, Data curation, Formal analysis, Funding acquisition, Investigation, Methodology, Project administration, Resources, Supervision, Validation, Visualization, Writing-original draft, Writing-review and editing;

